# Reduction of susceptibility to azoles and 5-fluorocytosine and growth acceleration in *Candida albicans* by glucose in urine

**DOI:** 10.1101/2021.05.10.443537

**Authors:** Yoshiki Hiyama, Toyotaka Sato, Satoshi Takahashi, Soh Yamamoto, Noriko Ogasawara, Naoya Masumori, Shin-ichi Yokota

## Abstract

*Candida species* are causal pathogens for urinary tract infections, vulvovaginitis, and balanitis. Diabetes mellitus is a risk factor for *Candida* infection. To investigate the potential effects of glucosuria on *Candida* spp. (*C. albicans, C. krusei*, and *C. glabrata*), we investigated the influence of their growth and antifungal susceptibilities by glucose in urine.

*Candida* spp. exhibited greater growth in urine with glucose (300 and 3,000 mg/dL) than in plain urine taken from healthy volunteers. After 24 h incubation, the viable cell number was more than 10-fold higher in the urine added 3,000 mg/dL glucose than in plain urine.

In antifungal susceptibility, more than 80% of *C. albicans* clinical isolates increased minimum inhibitory concentrations of azoles (fluconazole, itraconazole, voriconazole, and miconazole) and 5-fluorocytosine with the addition of glucose exceeding their breakpoints. This phenomenon was not observed in clinical isolates of *C. krusei* and *C. glabrata*. We observed the growth in the urine to which 3,000 mg/dL glucose was added even in the presence of a 128-fold higher minimum inhibitory concentration of fluconazole. In most of the *C. albicans* clinical isolates, the mRNA expression of the azole resistance genes *ERG11, CDR1, CDR2*, and *MDR1* increased in glucose-added urine compared with plain urine.

In conclusion, the growth of *C. albicans* is accelerated and azoles and 5-fluorocytosine become ineffective as a result of a high concentration of glucose in urine. These observations provide valuable information about the clinical course and therapeutic effects of azoles against *C. albicans* infections in patients with diabetes mellitus and hyperglucosuria.

**IMPORTANCE:** Diabetes mellitus is a chronic metabolic disease characterized by hyperglycemia and glucosuria, with a high risk of *Candida* infection. The current study demonstrated the acceleration of *Candida* growth and ineffectiveness of azoles and 5-fluorocytosine against *C. albicans* in urine in the presence of glucose. These observations provide novel and valuable information about the clinical course and antifungal treatment of *Candida* spp. in urinary tract and genital infections of diabetes mellitus patients. For the treatment of urinary tract infections caused by *Candida* spp., the guidelines do not mention glucosuria. Thus, this study suggests the necessity to conduct clinical evaluations for glucosuria in patients with diabetes mellitus who have urinary tract and genital infections with *Candida* spp.

## INTRODUCTION

Diabetes mellitus is a chronic metabolic disease characterized by hyperglycemia and glucosuria. Globally, an estimated 422 million people had diabetes worldwide (8.5% of the adult population) in 2014 with type 2 diabetes making up about 90% of the cases (1). Diabetes mellitus is a condition predisposing to infections. In the United States, 6 million diabetics were annually hospitalized, and 8-12% of them were admitted for infection management that was responsible for over 48 billion dollars in hospital charges (2).

Some of the cases are attributed to intrinsic infections by commensal bacteria or fungi (3). Epidemiological data on the association between diabetes and infection provide only limited scientific evidence that diabetes is associated with an increased risk of mortality (3, 4).

*Candida* spp. are commensals as a part of the normal human flora, and they are localized on the mucosae of the oral cavity, gastrointestinal tract and urogenital apparatus, and skin. In diabetes mellitus patients, as well as susceptible hosts such as elderly, hospitalized, and immunosuppressed patients, *Candida* spp. cause various infections such as oral candidiasis, peritonitis, vulvovaginitis, urinary tract infections, and balanitis (5). Their colonization increases in the oral cavity and gastrointestinal tract in patients with diabetes mellitus (6–8). *Candida albicans* is the most isolated fungal species in urinary tract infections, and in persons with diabetes mellitus with candiduria (the presence of *Candida* spp. in urine), there is a high risk of developing fungal urinary tract infections (8). It is believed that candiduria is one of the causal reasons for vulvovaginitis and balanitis (5). Therefore, detailed studies on candiduria and urinary *Candida* infections in diabetes mellitus would be important for control and treatment. However, the characteristics of *Candida* spp. in the glucosuria observed in patients with diabetes mellitus have not been well elucidated.

In this study, we investigated the effect of glucose addition to urine on the growth and susceptibility to antifungal agents of *Candida* spp. to simulate the glucosuria of diabetes mellitus.

## RESULTS

### Incidence of *Candida* spp. isolated from clinical glucosuria samples

We collected 553 urine specimens (527 and 26 were from patients without and with glucosuria, respectively) from patients without duplication (Table 1). Fungi were isolated from 9 (0.17%) specimens from patients without glucosuria, and all isolates were *Candida* spp. *Candida* spp. were also the solo fungi isolated from 4 (15.4%) specimens from patients with glucosuria. Fungi, namely *Candida* spp., were more frequently isolated from urine with glucosuria than from that without glucosuria (*p*<0.01).

**TABLE 1.** Microorganisms isolated from urine of patients positive and negative for glucosuria

| Glucosuria <sup>a</sup> | Bacteria |  | Fungi |
| --- | --- | --- | --- |
|  | Gram-negative | Gram-positive | *Candida* spp. <sup>b</sup> |
| Negative (527) | 299 (56.7%) | 354 (67.2%) | 9 (0.17%) |
| Positive (26) | 17 (65.4%) | 19 (73.1%) | 4 (15.4%)** |

a, Glucosuria was determined by using Meditape II 9U. Positive: >1,000 mg/dL of glucose.
b, included *C. albicans*, *C. glabrata*, *C. krusei*, and the *C. parapsilosis* group.
\*\**p* <0.01 vs. Negative.

### Growth of *C. albicans* in urine with various concentrations of glucose added

To determine the effect of glucose in urine, the growth rate of the *C. albicans*clinical isolates SMC2 and SMC40 was examined in glucose-added urine derived from healthy volunteers. Addition of more than 300 mg/mL glucose to urine markedly enhanced the growth (Fig. 1A and B). The viable colony count of *C. albicans* also exhibited enhancement of the growth rate at a glucose concentration in urine of more than 30 mg/dL (Fig. 1C). Enhancements of growth rates by the addition of glucose were also observed in urine derived from other three donors (Fig. 1D).

**FIG 1.**
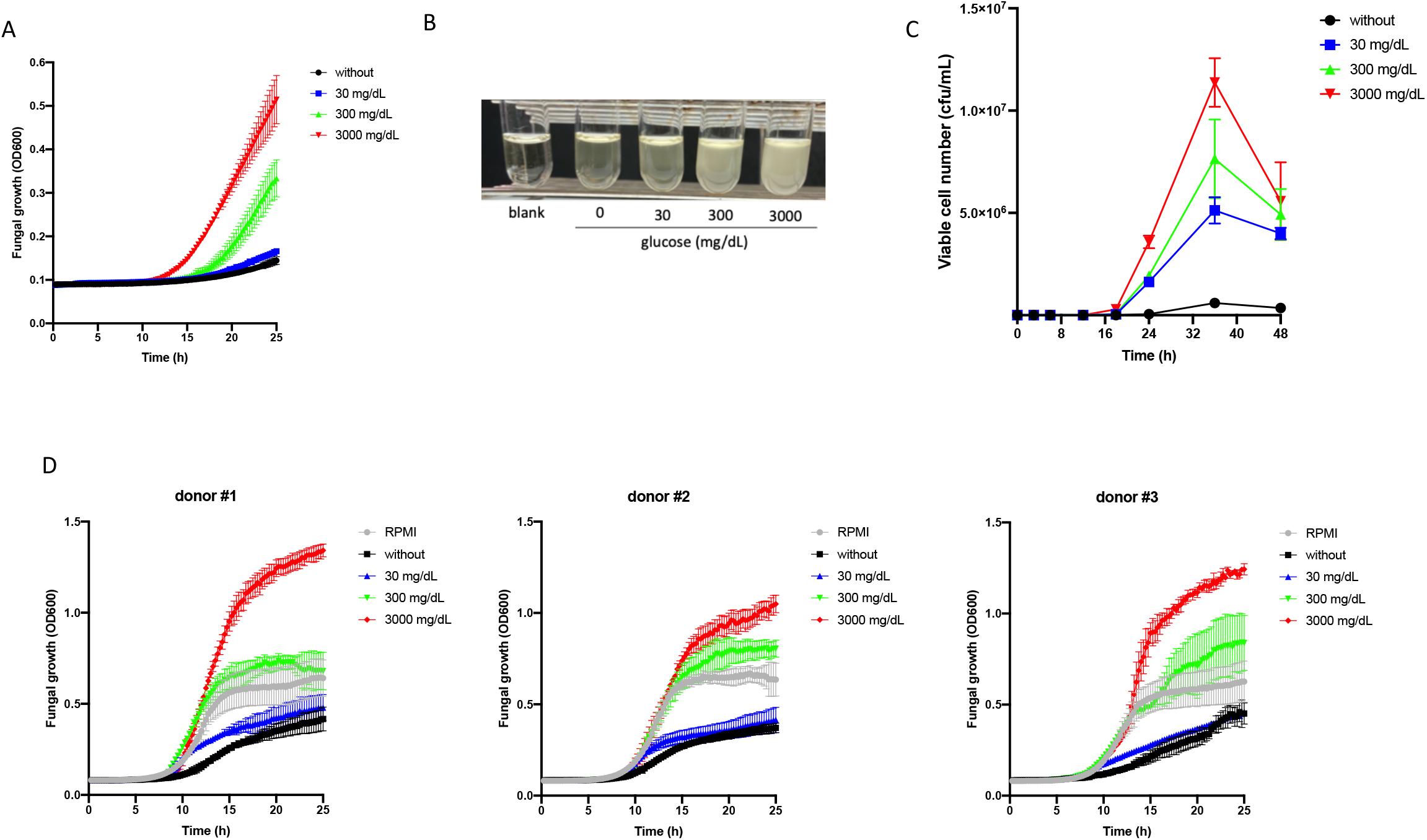
Growth rates of *C. albicans* in urine with various concentrations of glucose added. (A) Growth curves of *C. albicans* SMC2 in urine with glucose added were determined by measuring the values for the optical density at 600 nm. (B) The turbidity of *C. albicans* SMC40 culture at 48 h in urine with glucose added. (C) Growth determined by viable cell numbers (cfu/mL) of *C. albicans* SMC2 in urine with glucose. (D) Growth curves of *C. albicans* SMC40 culture in urine derived from three other donors, adding glucose, and RPMI 1640 medium.

We collected *Candida* spp. isolates from clinical urine specimens, and investigated the fungal growth for 36 *C. albicans* (other 14 *C. albicans* isolates that had no amplification of PCR products of seven housekeeping genes for MLST analysis were excluded from the evaluation.), 5 *C. krusei*, and 5 *C. glabrata* strains in urine. The 36 *C. albicans* clinical isolates exhibited genetic heterogeneity determined by nucleotide sequences obtained from MLST analysis (Fig. S1). All isolates of *Candida* spp., except one strain, exhibited markedly enhanced growth with the addition of glucose at 3,000 mg/dL in the urine compared to the absence of glucose addition (Fig. 2). After 24 h incubation, viable fungal numbers were more than 10-fold higher for *C. albicans* and *C. glabrata*, and more than 1,000-fold higher for *C. krusei* in urine in the presence of 3,000 mg/dL of glucose than in plain urine (Fig. 2).

**FIG 2.**
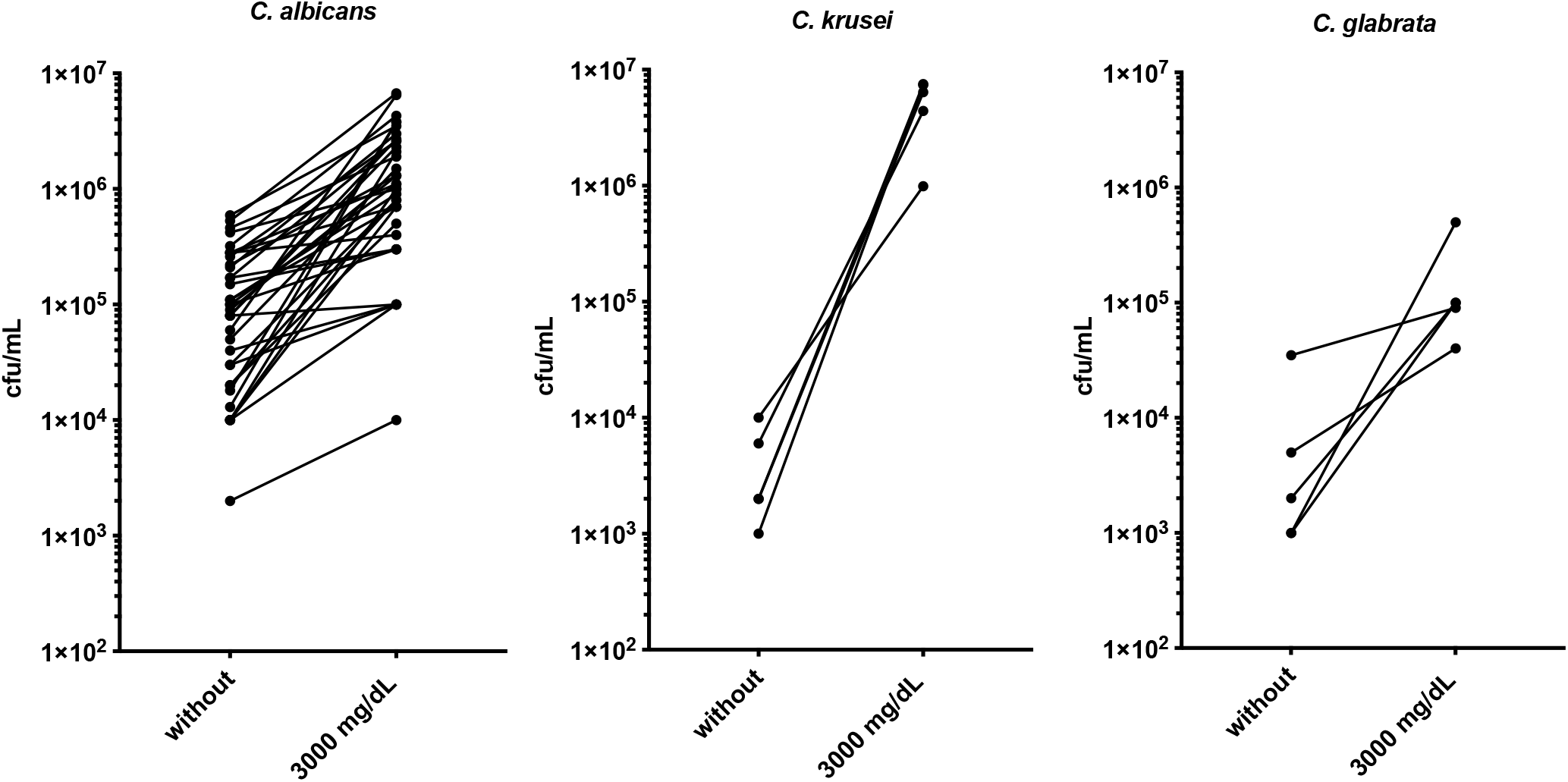
Viable cell numbers of clinical isolates of *C. albicans* (n=36), *C. krusei* (n=5), and *C. glabrata* (n=5) in urine with (3,000 mg/dL) or without addition of glucose after 24 h cultivation. Viable cell numbers were determined by cfu/mL.

### Antifungal susceptibility in urine with various concentrations of glucose

We measured minimum inhibitory concentrations (MICs) of various antifungal agents in urine with various concentrations of glucose (0, 300, and 3,000 mg/dL) for 50 *C. albicans* clinical isolates (Fig. 3). The MICs of all azole antifungal agents examined were similar in RPMI broth and urine without addition of glucose. The MICs of azoles (fluconazole, itraconazole, voriconazole, and miconazole), and 5-fluorocytosine were higher in urine in the presence of glucose than in plain urine (Fig. 3). The MIC levels were increased more than 128-fold for triazoles (fluconazole, itraconazole, and voriconazole) and 5-fluorocytosine, and 16-fold for miconazole. The phenotypes of fluconazole, itraconazole, and voriconazole were resistant. The resistance phenotypes of other antifungal agents (micafungin, caspofungin, and amphotericin B) were not altered in the glucose-added urine compared with RPMI broth and urine without addition of glucose. These phenomena were also observed using urine from other donors (Table S1).

**FIG 3.**
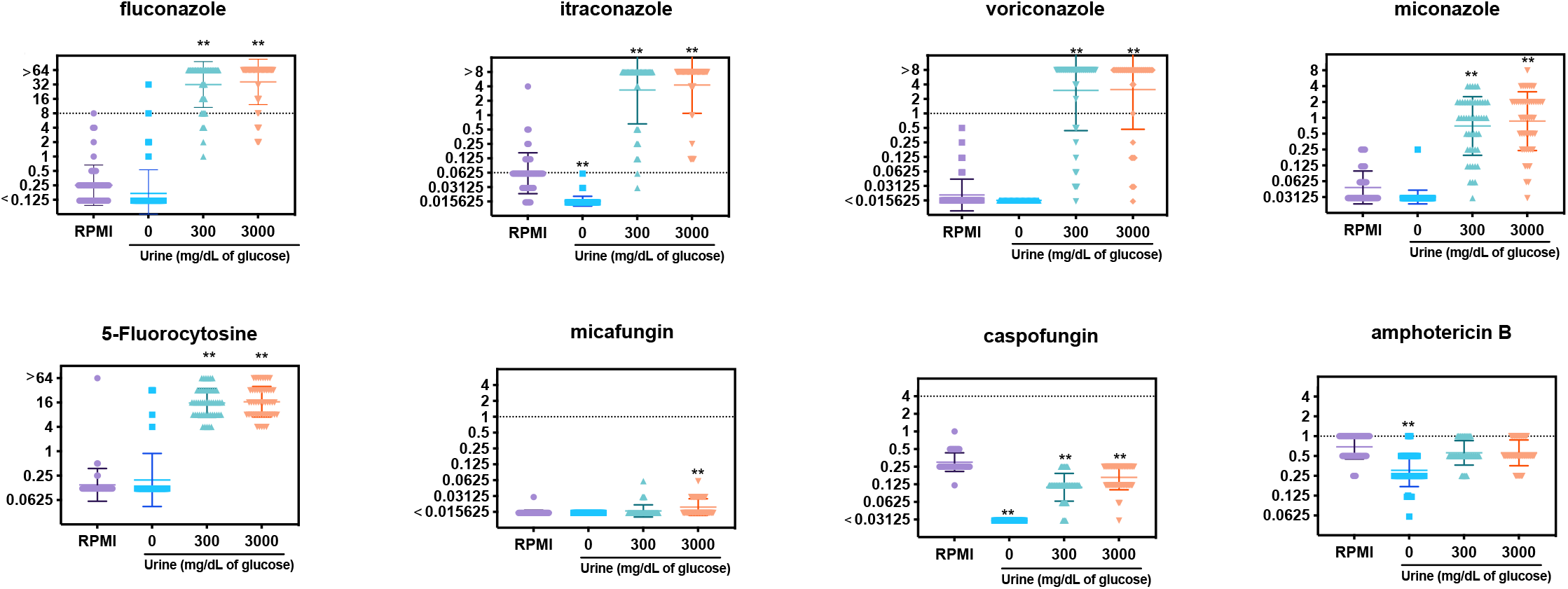
Minimal inhibitory concentrations (mg/L) of antifungal agents against *C. albicans* clinical isolates (n=50) cultured in RPMI1640 (standard procedure), urine, and urine added glucose at 300 and 3,000 mg/dL. *p*<0.01

The growth of *C. krusei* and *C. glabrata* in urine without addition of glucose was insufficient to determine MICs of the antifungals, except for amphotericin B (Tables S2 and S3). In most *C. krusei* and *C. glabrata* clinical isolates, the MICs of 5-fluorocytosine increased, and the MICs of caspofungin decreased in urine with 3,000 mg/dL glucose added compared with RPMI broth. These clinical isolates did not show apparent differences of the MICs of azoles, amphotericin B, and micafungin between RPMI broth and urine with 3,000 mg/dL glucose (Tables S2 and S3). The MICs of amphotericin B were similar or lower in urine with or without addition of glucose, than with RPMI broth.

### Time-kill assay in urine with addition of glucose

We evaluated the time-dependent antifungal effect by time-kill assay in urine using two *C. albicans* clinical isolates, SMC40 and SMC41 (Fig. 4). In the presence of amphotericin B, which is fungicidal, at a 2-fold higher MIC, complete fungicidal activity was observed within 24 h in RPMI broth, and in urine both with and without the addition of glucose (Fig. 4A and C). In the presence of fluconazole, which is fungistatic, at a 128-fold higher MIC, the viable cell numbers increased very slowly even after 2-day cultivation in RPMI broth and plain urine, and increased rapidly in urine with 3,000 mg/dL glucose (Fig. 4B and D). The increases in the glucose-added urine were significantly higher (*p*<0.05) than for RPMI broth and plain urine at 12 to 48 h.

**FIG 4.**
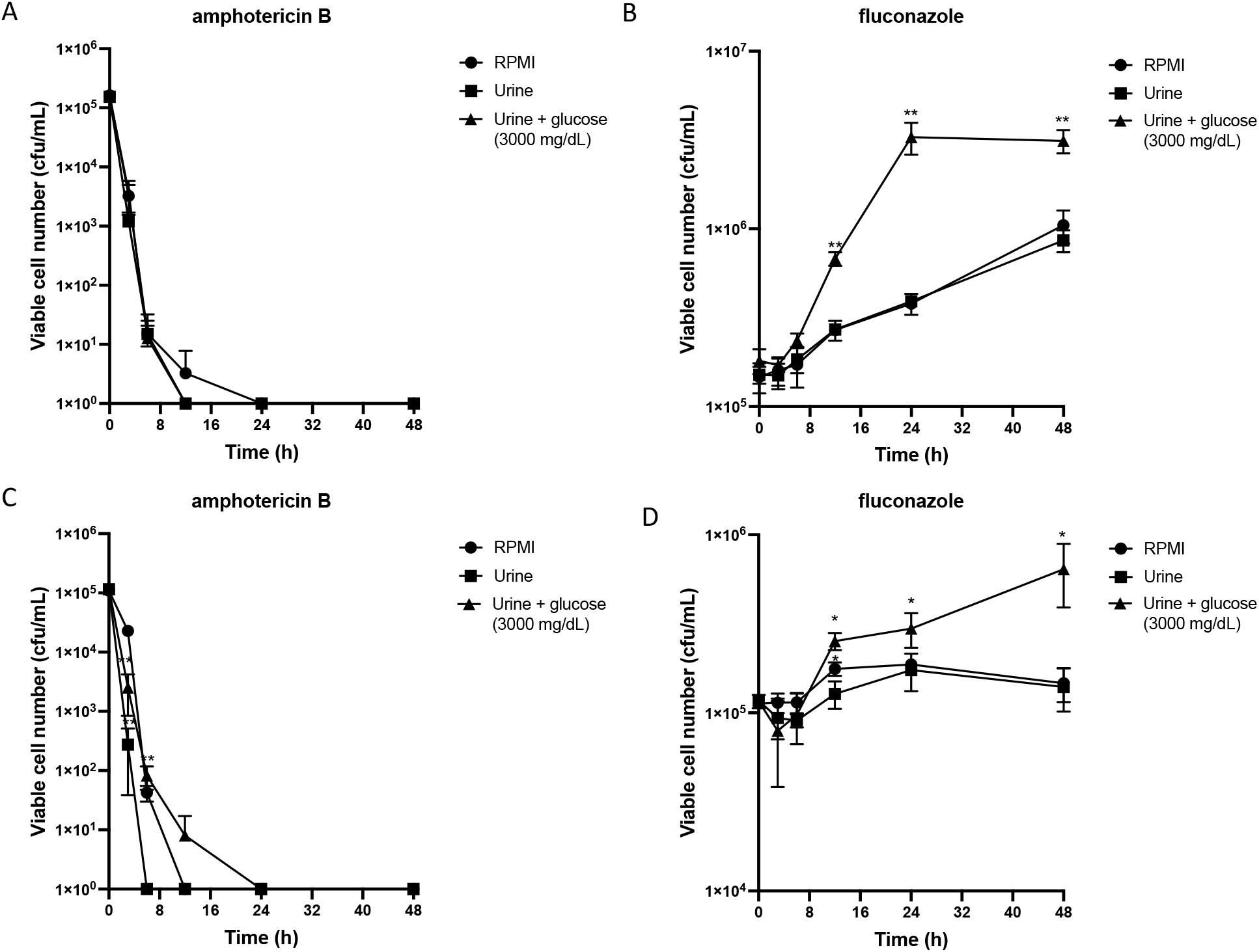
Time-kill assay of amphotericin B and fluconazole in *C. albicans* clinical isolates, SMC40 (A and B) and SMC41 (C and D) SMC 40 and SMC41 were cultured in RPMI 1640, urine, and urine to which 3,000 mg/mL glucose was added in the presence of amphotericin B or fluconazole. Concentrations of antifungals were 2-fold the MIC for amphotericin B (1 mg/L for both SMC40 and SMC41), and 128-fold higher the MIC for fluconazole (16 mg/L for SMC40 and 64 mg/mL for SMC41). The MICs were determined using RPMI 1640 medium. Viable cell numbers were determined as cfu/mL. *,*p*<0.05, **,*p*<0.01

### Expression of mRNA of azole-resistant-associated genes in glucose-added urine

We measured the expression of azole-resistance-associated genes (*ERG11, CDRI, CDR2*, and *MDR1*) in 36 *C. albicans* clinical isolates (Fig. 5). When compared with mRNA expression in the urine with and without addition of glucose, the levels were increased 3.1 ± 3.7-fold for *ERG11*, 2.5 ± 1.8-fold for *CDR1*, 8.5 ± 18.6-fold for *CDR2*, and 22.5 ± 34.5-fold for *MDR1* by addition of 3,000 mg/L glucose to urine.

**FIG 5.**
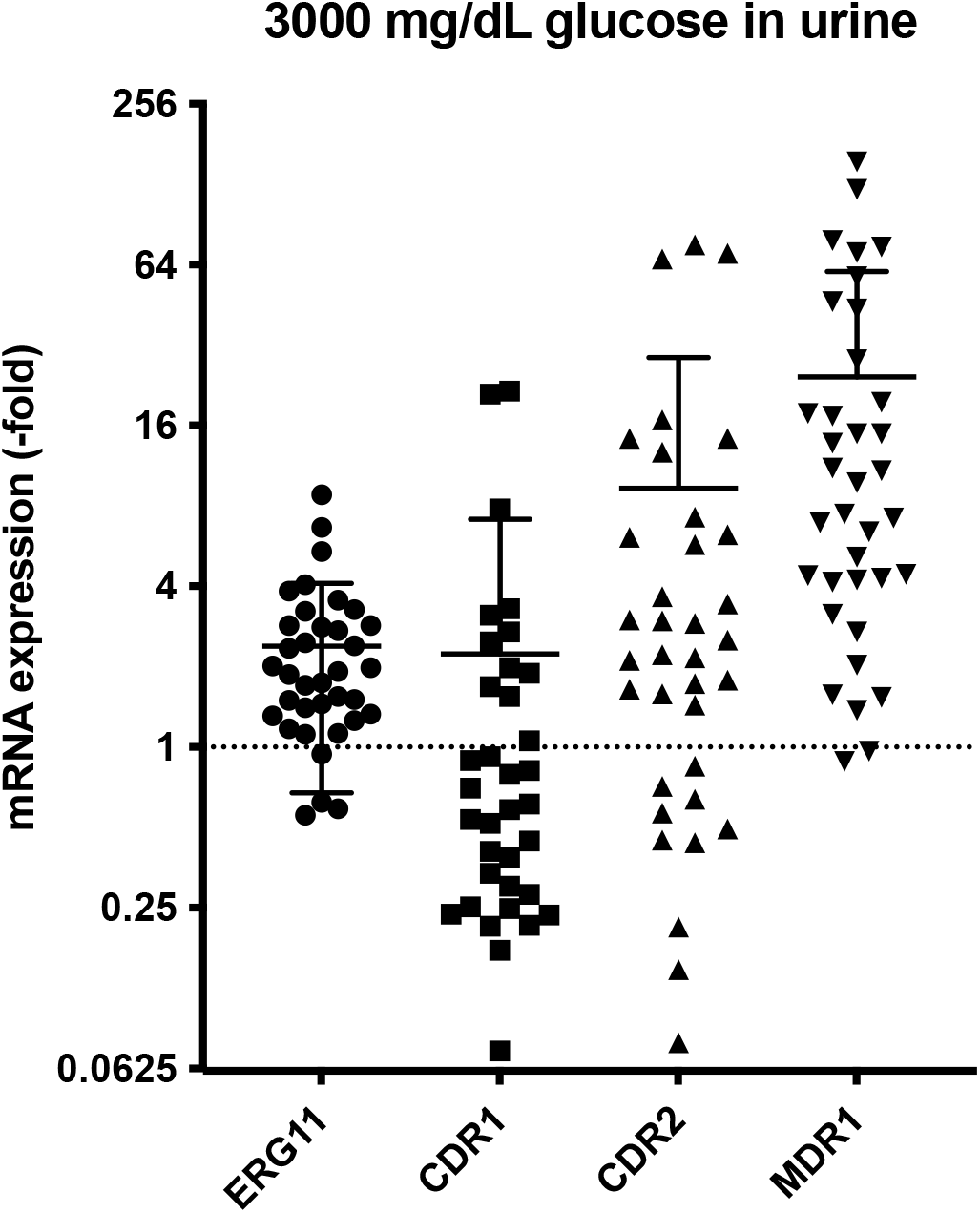
Effects of glucose on mRNA expression levels of *ERG11, CDR1, CDR2* and *MDR1* in *C. albicans* clinical isolates (n=36) during culture in urine and urine with 3,000 mg/dL glucose added. The expression levels were shown by relative expression level (-fold) compared with the levels of culture in plain urine.

### Susceptibility to antifungal agents of Mrr1-deleted *C. albicans* mutants

We measured MICs of antifungal agents in the azole-susceptible *C. albicans*strain SC5314 as the parent strain, and two derivatives of *MRR1*-deleted mutants (SCMRR1M4A and SCMRR1M4B) (Table 2). The mutants showed constitutive reduced expression of Mdr1 (9). In SC5314, the MICs of azoles and 5-fluorocytosine were markedly increased in the urine by addition of glucose, whereas those of candins and amphotericin B did not change. The MICs of azoles and 5-fluorocytosine against the two *MRR1*-deleted mutants were also increased by the addition of 300 mg/dL glucose; however, the increases MICs were smaller than those for SC5314. For instance, the MICs of fluconazole against SCMRR1M4A and SCMRR1M4B were 32- and 64-fold lower in urine with 300 mg/dL glucose than that against SC5314. The MICs of voriconazole against SCMRR1M4A and SCMRR1M4B were 2- and 8-fold lower (Table 2). Decreased MICs against the *MRR1*-deleted mutants compared to the parent strain were also seen in urine with 3,000 mg/dL glucose.

**TABLE 2.** Minimal inhibitory concentrations of antifungal agents against *MRR1*-deleted *C. albicans* mutants

| Antifungal<br>agents | SC5314 |  |  | SCMRR1M4A |  |  | SCMRR1M4B |  |  |
| --- | --- | --- | --- | --- | --- | --- | --- | --- | --- |
|  | (parent strain) |  |  | *(MRR1Δ::FRT/ MRR1Δ::FRT SC5314 clone A)* |  |  | *(MRR1Δ::FRT/ MRR1Δ::FRT SC5314 clone B)* |  |  |
|  | RPMI | Urine + 300 | Urine + 3000 | RPMI | Urine + 300 | Urine + 3000 | RPMI | Urine + 300 | Urine + 3000 |
| FLCZ | 0.25 | 64 | >64 | 0.25 | 1 | >64 | 0.5 | 2 | >64 |
| ITCZ | 0.06 | 8 | >8 | 0.06 | 0.12 | 2 | 0.12 | 0.12 | 4 |
| VRCZ | <0.015 | 0.25 | >8 | 0.03 | 0.03 | >8 | 0.03 | 0.12 | >8 |
| MCZ | <0.03 | 0.5 | 2 | 0.06 | 0.5 | 0.5 | 0.06 | 0.5 | 0.12 |
| 5-FC | <0.12 | 8 | 2 | <0.12 | 8 | 8 | <0.12 | 8 | 2 |
| MCFG | <0.015 | <0.015 | <0.015 | <0.015 | <0.015 | <0.015 | <0.015 | <0.015 | <0.015 |
| CPFG | 0.25 | 0.12 | 0.25 | 0.25 | 0.25 | 0.25 | 0.25 | 0.25 | 0.25 |
| AMPH-B | 0.5 | 0.25 | 0.25 | 0.5 | 0.25 | 0.25 | 0.5 | 0.25 | 0.5 |

4 Values are MICs (mg/L). FLCZ, fluconazole; ITCZ, itraconazole; VRCZ, voriconazole; MCZ, miconazole; 5-FC, 5-fluorocytosine; MCFG,
5 micafungin; CPFG, caspofungin; AMPH-B, amphotericin B.
6 Urine + 300, urine added 300mg/dL glucose; Urine + 3000, urine with 3,000mg/dL glucose added.

## DISCUSSION

We observed growth acceleration of *Candida* spp. in urine induced by addition of glucose in a concentration-dependent manner. This was consistent with a previous study (10); however, that study was conducted by using artificial urine and only one *C. albicans* strain. Our present study strengthened the previous observations in that: 1) the fungal growth acceleration was observed in natural urine derived from healthy donors as a result of adding glucose, and 2) this acceleration was observed in multiple *C. albicans*clinical isolates, in addition to *C. krusei* and *C. glabrata*. We believe that our study strengthens the evidence concerning the clinical risk of urinary tract infections caused by *Candida* spp. under glucosuria in diabetes mellitus.

For the treatment of type 2 diabetes mellitus, sodium glucose transporter 2 (SGLT2) inhibitors such as dapagliflozin, tofogliflozin, and empagliflozin have been widely used (11). SGLT2 mediates the tubular reabsorption of the majority of glomerular-filtered glucose. SGLT2 inhibitors suppress glucose reabsorption via an insulin-independent mechanism and thereby reduce blood glucose levels and increase urinary glucose excretion (12, 13). It is known that the glucose concentration in urine of patients administered SGLT2 reaches 3,000 mg/dL (14). We observed that the high glucose concentration in the urine conferred markedly enhanced fungal growth of *Candida* spp. A previous study reported a tendency towards an increased prevalence of *Candida* spp. in the urine of diabetic patients treated with canagliflozin (15). In addition, some other studies reported increased frequencies of urinary and genital infections, including vulvovaginal candidiasis and asymptomatic candiduria, during clinical trials of several SGLT2 inhibitors (15–19). Therefore, these cases might be attributable to fungal growth promotion in high glucosuria.

In the current study, we demonstrated dramatically decreased susceptibility to azoles, especially triazoles, and 5-fluorocytosine of *C. albicans* in glucose-added natural urine. This observation warrants great concern in the clinical setting because azoles are the most commonly used agents for the treatment of urinary tract infections caused by *C. albicans*, as well as its genital infections and vulvovaginal candidiasis. The ineffectiveness of azoles was observed in urine in the presence of glucose at a concentration of more than 300 mg/dL. This suggests high glucosuria in many of diabetes mellitus patients because 52.5% of the patients have glucose concentrations of more than 100 mg/dL in their urine regardless of SGLT2 inhibitor administration (20). The azole resistance phenotype in the presence of glucose in urine was observed in more than 80% of *C. albicans* clinical isolates. Therefore, high glucosuria might influence the risk of urinary and genital infections, and cause possible failure of antifungal treatment of *C. albicans* cases.

On the other hand, *Candida* spp. other than *C. albicans*, such as *C. krusei* and *C. glabrata*, also showed growth acceleration, but decreased susceptibility to antifungal agents with the addition of glucose to urine was not observed. The reason for the difference between *Candida* species is unknown. The biosynthesis and metabolism of the fungal cell membrane ergosterol, which is the target for azoles, is likely to be different among the various species. *C. albicans* can grow under 14α-sterol demethylase deficiency (21). On the other hand, *C. krusei* and *C. glabrata* are intolerant to the deficiency. The mechanisms of the differences, tolerance to 14α-sterol demethylase deficiency and decreasing susceptibility of azoles induced by glucose, between species need to be evaluated in the future.

To investigate the mechanism that confers the azole-resistant phenotype induced by glucose, we measured the gene expression of the azole resistance genes *ERG11*, *CDR1*, *CDR2*, and *MDR1*. Azoles contribute to antifungal activity by inhibiting 14α-sterol demethylase encoded by *ERG11*, which is involved in the biosynthesis of ergosterol (22). Increased expression of Erg11 overcomes the activity of azoles and thereby increases the azole resistance level (22). Cdr1, Cdr2, and Mdr1 are efflux transporters that excrete multiple compounds, including azoles. Overexpression of these genes confers azole resistance (22). We found that *C. albicans* clinical isolates had enhanced expression levels of *ERG11, CDR1, CDR2*, and *MDR1* in urine to which glucose was added. *CDR2* and *MDR1* expression in particular increased extremely (Fig. 5). To confirm the contribution of Mdr1 to the azole resistance in urine, we performed antifungal susceptibility tests using *MRR1*-deleted *C. albicans* mutants. Mrr1 is a regulator of Mdr1, and mutations in *MRR1*result in constitutively reduced expression of Mrd1 (23). The enhancement of azole resistance levels induced by glucose in urine was partially inhibited in the *MRR1*-deleted mutants compared with the parent strain (Table 2). Thus, the azole resistance induced by glucose in urine might be partly contributed to by Mrr1-dependent overexpression of the azole resistance genes.

In conclusion, growth acceleration in *Candida* spp. and ineffectiveness of azoles and 5-fluorocytosine against *C. albicans* occur in urine in the presence of glucose at high concentrations comparable to glucose concentrations in the urine of diabetes mellitus patients, especially during treatment with SGLT2 inhibitors. The current results provide novel and valuable information about the clinical course and antifungal treatment of *Candida* spp. in urinary tract and genital infections. For the treatment of urinary tract infections caused by *Candida* spp., the guidelines do not mention glucosuria (24). We thus need to conduct clinical evaluations for glucosuria in patients with diabetes mellitus who have urinary tract and genital infections with *Candida* spp.

## Material and Methods

### Collection of urine specimens and isolation of bacteria and fungi

We collected the urine specimens from daily laboratory diagnostic tests in the Department of Urology, Sapporo Medical University Hospital (Sapporo, Japan) in 2017. This study was approved by the Sapporo Medical University Ethics Committee (No. 302-1031). We collected urine specimens without glucosuria and with glucosuria (with a glucose concentration of more than 1,000 mg/dL) determined by using Meditape II 9U (Sysmex, Tokyo, Japan). The identification of bacteria and fungi obtained by cultivation tests was performed using a MALDI Biotyper (Bruker, Billerica, MA). Only the first culture of one episode was investigated to avoid duplication. We calculated the proportions of gram-negative and gram-positive bacteria, and fungi in samples with and without glucosuria.

Urine used for fungal culture experiments was obtained from healthy volunteers, and filtrated with a 0.45 μm-pore membrane (TPP Filtermax, Merk KGaA, Darmstadt, Germany).

### Isolation and characterization of *Candida* spp. strains

We collected clinical isolates of *Candida* spp. (50 *C. albicans*, 5 *C. krusei*, and 5 *C. glabrata* isolates), and used them for the experiments. These isolates were isolated from patients’ urine at Sapporo Medical University Hospital in 2019-2020. The identification of bacteria and fungi was performed using a MALDI Biotyper. Multilocus sequence typing (MLST) of *C. albicans* based on seven housekeeping genes (*AAT1a*, *ACC1, ADP1, MP1b, SYA1, VPS13*, and *ZWF1b*) was conducted as previously described (25, 26), to confirm genetic heterogeneity of these isolates. A phylogenetic tree based on the nucleotide sequence data of MLST analysis for the *C. albicans* isolates was constructed based on the neighbor-joining method (27) using MEGA7 (28).

*C. albicans* SC5314, SCMRR1M4A and SCMRR1M4B were kindly provided by Dr. Joachim Morschhäuser (9). SCMRR1M4A and SCMRR1M4B are homozygous *MRR1Δ* mutants that share decreased *MDR1* promoter activity originated from SC5314 (9).

### Growth curves of *C. albicans* in urine with various concentrations of glucose

*C. albicans* strains, SMC2 and SMC40, were cultured on Sabouraud agar plates (Nissui Pharmaceutical, Tokyo, Japan) for 24-48 h at 37°C. One colony was picked and inoculated into 100 μL of urine with or without glucose in 96-well plates (VIOLAMO, Osaka, Japan). The plates were cultivated at 37°C with shaking at 180 rpm. Growth curves were determined by measuring values for OD_600_ at every 15 min for 25 h by using an Infinite M200 PRO multimode microplate reader (Tecan, Männedorf, Switzerland). Growth curves were obtained from the average of quadruple experiments.

To determine the viable cell number in urine culture with or without addition of glucose, colony formation units (cfu) were measured. *C. albicans* at 100 cfu/mL was inoculated into the urine obtained from healthy volunteers with or without glucose (30, 300, and 3,000 mg/dL). The urine was cultured at 37°C with shaking at 180 rpm for 1, 3, 6, 12, 24, 36, and 48 h. After the cultivation, series of dilutions with 0.85% NaCl were spread on Sabouraud agar plates, and incubated for 24 h at 37°C, after which formed colonies were counted.

### Antifungal susceptibility

MICs of antifungal agents (fluconazole, itraconazole, voriconazole, miconazole, micafungin, caspofungin, amphotericin B, and 5-fluorocytosine) were determined by the broth microdilution method according to Clinical and Laboratory Standards Institute (CLSI) guidelines (29) using a susceptibility test kit (Eiken Chemical, Tokyo, Japan) according to the manufacturer’s instructions. RPMI 1640 (RPMI) broth (Eiken Chemical) was used as a medium. After 24 or 48 h, the growth in each well of a 96-well plate was measured by the OD_630_ value using an Infinite M200 PRO multimode microplate reader. For all antifungal agents except amphotericin B, a value of less than the IC_50_ [50% growth inhibition of the well of a positive control that was without any agents in RPMI broth] was defined as the MIC. The breakpoints of itraconazole and amphotericin B were according to EUCAST (30) because there is no definition in the CLSI guidelines.

### Time-kill assay

Fluconazole and amphotericin B were purchased from FUJIFILM Wako Pure Chemicals (Osaka, Japan). *C. albicans* SMC40 and SMC41 were grown overnight at 37°C on tryptic soy broth (TSB). Cells of each strain were added at a concentration of 10^5^ cfu/mL to urine with or without addition of glucose and an antifungal agent. The concentrations of glucose were 300 and 3000 mg/dL. The concentrations of fluconazole were 16 and 64 μg/mL, which were 128-fold higher concentrations of fluconazole MICs against SMC40 and SMC41, respectively. The concentration of amphotericin B was 1 μg/mL, which was 2-fold higher than the concentration of amphotericin B MICs against both strains. The urine was incubated with shaking at 37°C. Aliquots of urine collected at 1, 3, 6, 24, and 48 h were inoculated and cultured on a Sabouraud agar plate to determine the viable cell numbers (cfu/mL).

### Reverse transcription-quantitative PCR (RT-qPCR)

Overnight cultures of *C. albicans* clinical isolates in TSB were diluted 1:25 in RPMI or urine with 3000 mg/dL glucose added, and then cultured for 3 h at 37°C. RNA was isolated using Yeast Processing Reagent (Takara Bio, Shiga, Japan) and RNeasy Plus mini kit (Qiagen, Hilden, Germany) according to the manufacturers’ instructions.

The concentration of RNA was measured spectrophotometrically using an Infinite M200 PRO. RNA (0.5 μg) was used to synthesize cDNA by utilizing ReverTra Ace reverse transcription-quantitative PCR master mix with genomic DNA (gDNA) remover (Toyobo, Tokyo, Japan). Expression of *ERG11*, *CDR1*, *CDR2* and *MDR1* was determined as described previously (31) using KOD SYBR qPCR mix (Toyobo). The PCR cycling conditions were as follows: initial activation at 95°C for 5 min, followed by 40 cycles at 95°C for 10 s and 55°C for 30 s. Reactions were performed in a LightCycler 480 II (Roche, Mannheim, Germany). The *ACT1* gene served as an endogenous reference for normalizing expression levels. All primer sets were used as previously described (31). The change in fold expression in urine with 3,000 mg/dL glucose added (vs. plain urine) was calculated by the ΔΔCT method. Data are expressed as the means ± standard deviations from three independent experiments.

### Statistical analysis

Significant differences were determined using Fisher’s exact test (Table 1), the Kruskal-Wallis (Fig. 3), and Wilcoxon signed-rank test (Fig. 4). *p*<0.05 was considered significant.

## Authorship

All authors meet the ICMJE authorship criteria.

## ACKNOWLEGEMENTS

We thank Dr. Joachim Morschhäuser for kind provision of *MRR1*-deletd *C. albicans* mutants. This work was supported by grants from the Japan Agency for Medical Research and Development (AMED) (JP20ak0101118h0002). This work was also partly supported by a grant from JSPS KAKENHI (19K18566, JP19K16648, and JP20H03488). The funding sources did not play any role in the study design; in the collection, analysis and interpretation of data; in the writing of the report; and in the decision to submit the article for publication.

## Supplemental data

**Table S1. Minimal inhibitory concentrations of antifungal agents against *C. albicans* SMC40 in RPMI1640 (standard procedure) and urine derived from different donors in the presence (300 and 3,000 mg/mL) and absence of glucose (Excel file)**

**Table S2. Minimal inhibitory concentrations of antifungal agents against *C. krusei*isolates in RPMI1640 (standard procedure) and urine in the presence (3,000 mg/mL) and absence of glucose (Excel file)**

**Table S3. Minimal inhibitory concentrations of antifungal agents against *C. glabrata* isolates in RPMI1640 (standard procedure) and urine in the presence (3,000 mg/mL) and absence of glucose (Excel file)**

**FIG S1. Phylogenetic tree based on MLST analysis in *C. albicans* clinical isolates (n=36)**

